# Geographical distribution, disease association and diversity of *Klebsiella pneumoniae* KL and O antigens in India: roadmap for vaccine development

**DOI:** 10.1101/2024.03.15.585133

**Authors:** Varun Shamanna, Srikanth Srinivas, Natacha Couto, Geetha Nagaraj, Shyama Prasad Sajankila, Harshitha Gangaiah Krishnappa, Kavitha Arakalgud Kumar, David Aanensen, Kadahalli Lingegowda Ravikumar, GHRU India Consortium

**Author notes:** **Corresponding Author:** Varun Shamanna. **Repositories:** All the WGS data has been submitted to ENA under the Bioproject numbers PRJEB29740 and PRJEB50614. All the R scripts used in the study have been deposited in Figshare (https://doi.org/10.6084/m9.figshare.25414807.v1).

## Abstract

*Klebsiella pneumoniae* poses a significant healthcare challenge due to its multidrug resistance and diverse serotype landscape. This study aimed to explore the serotype diversity of 1072 *K. pneumoniae* and its association with geographical distribution, disease severity and antimicrobial/virulence patterns in India. Whole-genome sequencing was performed on the Illumina platform, and genomic analysis was carried out using the Kleborate tool. KL64 (n=264/1072, 26%), KL51 (249/1072, 24%), KL2 (n=88/1072, 8%), O1/O2v1 (n=471/1072, 44%), O1/O2v2 (n=353/1072, 33%), and OL101 (n=66/1072, 6%) were the most prevalent serotypes. The study identified 119 different sequence types (STs) with varying serotypes, with KL64 being the most predominant in ST231 (26%). O serotypes were strongly linked with STs, with O1/O2v1 predominantly associated with ST231 (44%). Simpson’s diversity index and Fisher’s exact test revealed higher serotype diversity in the north and east regions, along with intriguing associations between specific serotypes and resistance profiles. No significant association between KL or O types and disease severity was observed. Furthermore, we found no specific association of virulence factors with KL types or O antigen types (P>0.05). Conventionally described hypervirulent clones (i.e., KL1 and KL2) in India lacked typical virulent markers (i.e., aerobactin), contrasting with other regional serotypes. The cumulative distribution of KL and O serotypes suggests that future vaccines may have to include either ∼20 KL types or 4 O types to cover >85% of the carbapenemase-producing Indian K. pneumoniae population. The findings underscore the need for a vaccine with broad coverage to address the diverse landscape of K. pneumoniae strains in different regions of India. Understanding regional serotype dynamics is pivotal for targeted surveillance, interventions, and tailored vaccine strategies to tackle the diverse landscape of *K. pneumoniae* infections across India.

**Data Summary:**

- All the sequenced data has been submitted to the European Nucleotide Archive (ENA) under the Bioproject numbers PRJEB29740 and PRJEB50614. Run Accessions and Biosample numbers are provided in Supplementary Table 1 with corresponding metadata for each sample used in the study.
- The Microreact link for the genomic analysis is provided (https://microreact.org/project/oqKM84GBszEPW9Emt2FKnP-klebsiella-pneumoniae-indian-serotypes).
- The pipelines used in the study are published in gitlab (https://gitlab.com/cgps/ghru/pipelines).
- The tools’ details and the implementation of the pipelines are described in protocols.io (https://www.protocols.io/view/ghru-genomic-surveillance-of-antimicrobial-resista-bp2l6b11kgqe/v4).
- The R scripts used with all the input files used for each script have been published in Fishare (https://doi.org/10.6084/m9.figshare.25414807.v1)

**Impact Statement:** *Klebsiella pneumoniae* produces polysaccharide capsules, which serve as both epidemiological markers and significant virulence factors. The increasing accessibility of whole genome sequencing has made it easier than ever to investigate this capsule diversity. This study is the first of its kind in India to comprehensively investigate the serotype diversity of *K. pneumoniae* strains and their association with disease severity, antimicrobial resistance/virulence patterns, and geographical distribution across various regions of the subcontinent. This multi-dimensional analysis not only provides valuable insights into the molecular epidemiology of *K. pneumoniae* in India but also offers crucial data for the development of targeted interventions, including vaccine formulations tailored to address the prevailing serotypes. These findings serve as a foundation for informed decision-making in the management and prevention of *K. pneumoniae* infections, ultimately contributing to improved public health outcomes in the region.

## Introduction

*Klebsiella pneumoniae* (*K. pneumoniae*) is a Gram-negative bacterium that is a major cause of nosocomial infections [1], including pneumonia, sepsis, and urinary tract infections [2]; [3]. Its increasing resistance to antimicrobials poses a serious public health threat [4]. Genome-based surveillance is urgently needed to control the emerging threat of *K. pneumoniae [5]*. Recent advances in understanding the population structure of *K. pneumoniae* have revealed an immense genomic diversity, providing a framework for pathogen tracking [6]; [7]. The emergence of multidrug-resistant (MDR) *K. pneumoniae* strains, resistant to multiple antimicrobials, is a significant concern, especially in countries like India, where treatment becomes challenging. [8]; [9]; [10]; [11]. MDR hypervirulent *K. pneumoniae* (hvKp) strains have been emerging, spawning a new generation of hypervirulent “superbugs” [12]. Newer therapeutics, such as vaccines or monoclonal antibodies, are needed to combat MDR *K. pneumoniae*. Vaccines could help to prevent infections from occurring in the first place and/or they could help to reduce the severity of infections in affected individuals. Human monoclonal antibodies, on the other hand, can rapidly progress to innovative prophylactic and therapeutic solutions (Roscioli et al., 2024).

Capsular polysaccharides (CPS, encoded by the K locus) and lipopolysaccharides (LPS, encoded by the O locus) are major virulence factors of *K. pneumoniae*, and they are responsible for protecting the bacterium from the host’s immune system. Bacterial capsule-targeted vaccines, such as those designed for *Streptococcus pneumoniae, Neisseria meningitidis*, and *Haemophilus influenzae*, have demonstrated significant effectiveness in preventing illnesses caused by these encapsulated pathogens [13]. Currently, efforts are in place to develop vaccines or monoclonal antibodies against *K. pneumoniae*. The increased immunogenicity and enhanced surface exposure of CPS and LPS in *K. pneumoniae* render them appealing candidates [13–15]. A bioconjugation approach based on glycoengineered *Escherichia coli* expressing *K. pneumoniae* KL1 and KL2 antigens led to the production of IgG against both glycans in mice conferring protection against lethal challenges with KL1 and KL2 strains [16]. Affinivax is working on an LPS-based formulation that combines antigens O1, O2, O3, and O5 with the type III fimbriae adhesion MrkA to generate a multiple-antigen presenting system (MAPS) [17]. However, the wide range of CPS and LPS types makes it challenging to provide comprehensive coverage [18]. The structural variability, defective capsule or O-antigen production, and variations in the geographic distribution of serotypes limit the potential coverage of CPS/LPS-based vaccines or monoclonal antibodies [19].

There are over 80 capsular serotypes (encoded by the K-loci) of *K. pneumoniae [20]*; [21], and the prevalence of these serotypes varies from region to region. Understanding the distribution of *K. pneumoniae* serotypes is important for vaccine and monoclonal antibodies development, as a vaccine targeting the most prevalent serotypes in a given region is more likely to be effective. Among the various capsular (KL) types, KL1 and KL2 are often linked to high virulence, and more concerningly, isolates with KL47 and KL64 are often linked to both hypervirulence and carbapenem resistance [22], which present significant challenges for antimicrobial therapy [23,24]. These isolates are referred to as hypervirulent carbapenem-resistant *K. pneumoniae* (hv-CRKP) emphasizing the urgent need to design and develop broad-spectrum therapeutic drugs or vaccines against *K. pneumoniae* isolates of serotypes KL1, KL2, KL47, and KL64 [25]. In India, there are few studies with a limited number of samples. These have reported KL51 and KL64 as the most common *K. pneumoniae* serotypes, which are associated with specific sequence types (STs), ST231-KL51 and ST147-KL64 [26]; [27]. The frequency of other KL types varies in different studies based on the sample sizes. The globally prevalent, hypervirulent serotypes KL1, KL2, and KL20 were rarely found in these Indian studies.

The O-antigen is a significant virulence factor for *K. pneumoniae*. It aids the bacterium in evading the host’s immune system and attaching to host cells. The various O-antigen serogroups have different antigenic properties, which can affect the bacterium’s ability to cause disease. The O-antigen of *K. pneumoniae* can be divided into different groups based on their unique structures and antigenic properties. *K. pneumoniae* has eleven characterised LPS serogroups. Just four serogroups: O1, O2a, O3, and O5 are expressed by over 80% of all isolates [13]; [15]. In India, O1 and O2 are the most prevalent types, collectively constituting over 70% of the isolates, as indicated in prior studies [28]; [10]; [26]. However, it is worth mentioning that the number of studies conducted in the Indian setting is limited and these studies represent smaller-scale investigations with relatively modest sample sizes, underlining the need for more comprehensive research in this context.

Understanding the prevalence of *K. pneumoniae* serotypes in India will help the design of an effective vaccine that covers the main circulating lineages in the country. For that reason, in this study, we aim to identify the K and O loci of a large collection of country-wide *K. pneumoniae* strains causing infection in India. Furthermore, our study explores the relationship between these serotypes and disease severity, geographic distribution, and virulence and antimicrobial resistance (AMR) characteristics, which adds to our understanding of their intricate interactions in the Indian setting. This multi-pronged approach will help India to develop effective and specialised vaccines against *K. pneumoniae*.

## Materials and Methods

### Bacterial Isolates and Phenotypic Characterization

The bacterial isolates used for this study comprised of 1072 putative *K. pneumoniae* isolates primarily sourced from hospital infections and obtained from the years 2014 to 2022 across India. This includes 307 isolates from our previous study [26]. The phenotypic characterization was done at the Central Research Laboratory, Kempegowda Institute of Medical Sciences (KIMS) using the VITEK 2 (bioMérieux, Marcy-l’Étoile, France) compact system. The Ethical approval for the study was obtained from the KIMS ethical committee with the study number KIMS/IEC/27/2017. The strain details are provided in Supplementary Table 1.

### Sequencing and Genomic Analyses

#### Whole-genome sequencing, assembly, and annotation

Genomic DNA was extracted and isolated from the bacterial isolates using the QIAamp DNA mini kit (Qiagen, Hilden, Germany) and quantified using the Qubit double-stranded DNA kit (ThermoScientific, Massachusetts, United States) as instructed by the manufacturer. Double-stranded DNA libraries with 450 bp insert size were prepared using the ultraFS-II kit (New England Biolabs, London, United Kingdom). The QC check for the prepared libraries was done on an Agilent Tapestation (Santa Clara, California, USA) and libraries were sequenced on the Illumina MiSeq platform (Illumina, San Diego, California, United States of America) with paired-end reads of 250 bp length. All the WGS data generated as a part of this study were submitted to the European Nucleotide Archive (ENA) under the Bioproject numbers PRJEB29740 and PRJEB50614 with accession IDs provided in Supplementary Table 1.

The bioinformatic analysis was conducted using Nextflow pipelines created as part of the Genomic Surveillance of Antimicrobial Resistance-AMR project available at protocols.io [29]. The pipeline performs assembly using SPApades assembler v3.14 [30]. Quality control of sequence data was evaluated for the following parameters: (i) the basic statistics of raw reads, (ii) the assembly statistics, (iii) contamination due to single nucleotide variants (SNV) and sequences from different species, (iv) Species prediction using Bactinspector and (v) Overall QC as Pass, Warning or Fail for each isolate based on these different parameters as described in the pipeline. All the quality metrics were combined using Multiqc and Qualifyr to generate web-based reports [31]. The passed assemblies were annotated with Prokka v1.5 [32].

#### *In silico* genomic characterization

Kleborate v2.3.2 (https://github.com/katholt/Kleborate) is a designated genotyping tool developed for *Klebsiella* spp. It integrates multiple analysis steps, including multi locus sequence typing (MLST), and identification of virulence and acquired resistance genes, to provide a comprehensive genotypic profile of the isolates [33]. KL and O antigens were identified using Kaptive [34].

#### Variant detection and phylogenetic analysis

Genome mapping of the 1072 isolates to the reference genome of *K. pneumoniae* (strain NTUH-K2044, GCF_009497695.1) was done using the GHRU-SNP phylogeny pipeline v1.2.2 (https://gitlab.com/cgps/ghru/pipelines/snp_phylogeny). The mobile genetic elements (MGEs) were masked in the pseudo genome alignment using MGEmasker [35], and the recombinant regions of the genome were removed using the Gubbins algorithm v2.0.0 [36]. A maximum-likelihood tree was built utilising the non-recombinant SNPs using IQ-tree [37] with 100 bootstrap replicates and parameters -czb to collapse near-zero branches, and a general time-reversible (GTR) model. Phylogeographic analysis and visualisation were performed on Microreact [38].

#### Statistical Analysis and Plots

We used the Rstudio server v2023.03.0 Build 386 for descriptive and statistical analysis. Plots were generated using the ggplot package [39] and for data interpretation, the dplyr [40] and tidyr [41] packages were used. All the code used in the study is published on Zenodo and the link is provided.

## Results

### Overview of the collection

The collection included 1072 isolates from 38 different hospitals located in 19 different states across India from 60% (646/1072) male and 40% (426/1072) female population. Of the 1072 isolates, 65 (1%) isolates were obtained from children <24 months of age, and overall patient age ranged from 1 to 96 years with a median age of 48 years. The geographical representation and the timeline of the samples are shown in Supplementary Figure 1 and also provided in Supplementary Table 1.

The isolates were classified into two categories: invasive and non-invasive based on the site of collection and specimen type. Among the invasive specimen types, blood samples emerged as the predominant source, yielding 217 *K. pneumoniae* isolates followed by endotracheal aspirates (ETA) with 147 isolates and body fluids with 50 isolates. In the non-invasive specimen types, urine samples were predominant, with 257 isolates. Pus specimens were followed by 154 isolates and sputum samples with 140 isolates. It is noteworthy that 90% of the *K. pneumoniae* isolates were limited to 6 sample types and were not commonly found from other sources. The complete distribution of different specimens is represented in Figure 1.

**Figure 1.**
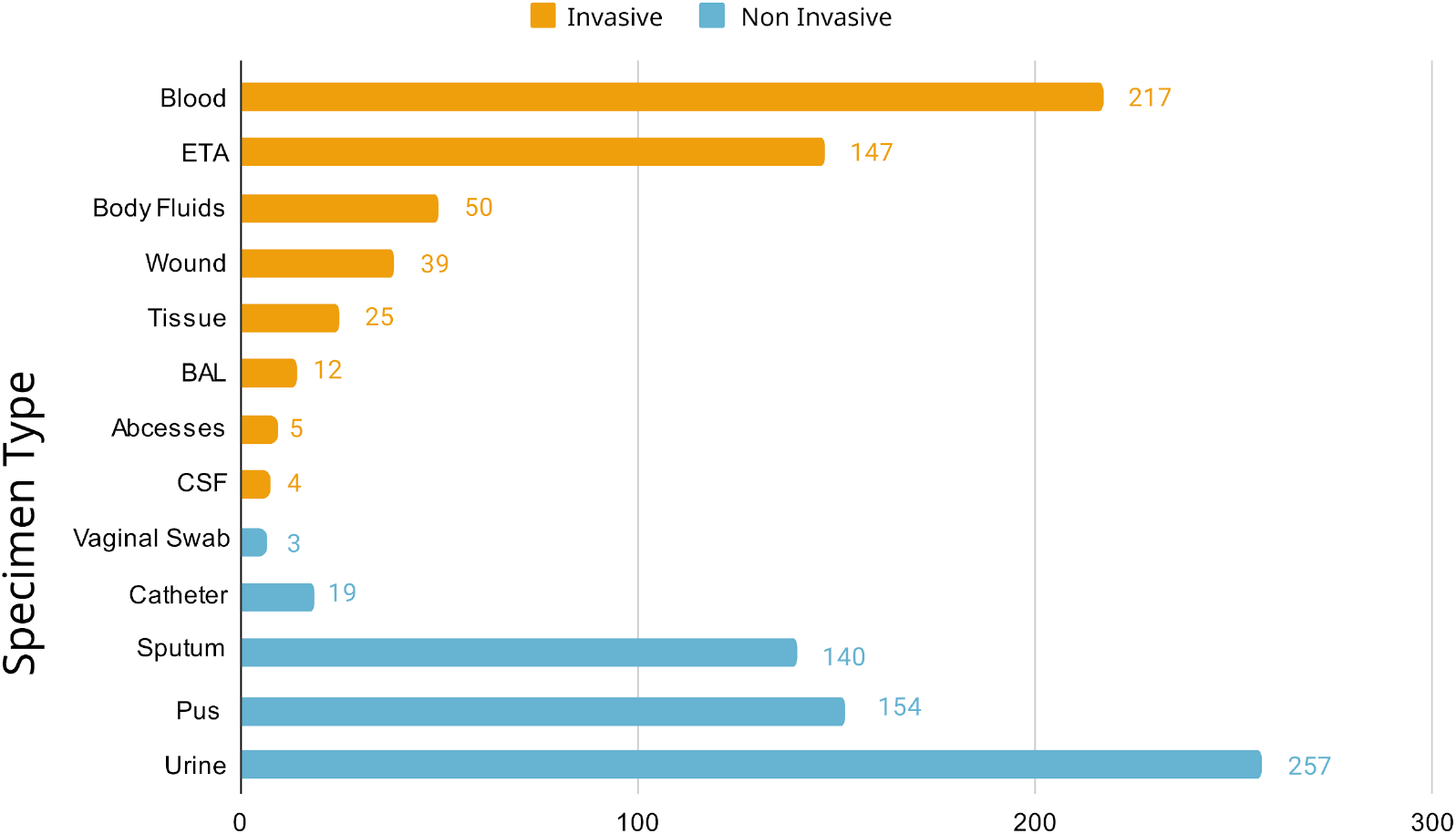
Distribution of specimen types, classified as invasive and non-invasive, from which *Klebsiella pneumoniae* was recovered.

### K and O loci Diversity

In our collection of non-redundant genomes, we identified 79 distinct KL. The most prevalent K-loci were KL64 (n = 274), KL51 (n = 249), KL2 (n = 88), and KL10 (n = 50). The top three K-loci contributed to 59% of the isolates. The top 10 major KL types are provided in Table 1 and the complete distribution of each K-loci is provided in Supplementary Table 2. The major KL types in both invasive and non-invasive specimen sources were similar, i.e. KL64 was a major KL in invasive sources (n=122) and also in non-invasive sources (n= 152) (Figure 2) (Suppl. Table 2). Hence, we assessed the association between KL sample types (invasive and non-invasive) using Fisher’s exact test [42]. The most prevalent K-loci, namely KL64 (P-value: 0.441), KL51 (P-value: 0.469), and KL2 (P-value: 0.266) were not statistically associated either with invasive or non-invasive disease. Only KL10 was significantly associated with non-invasive disease (P-value: 0.041), indicating a potential link.

**Table 1.** KL distribution among invasive and non-invasive specimen types. The top 10 KL of frequency >15 are shown individually and the lesser predominant KL <15 counts are grouped. The top 10 STs are shown individually and others are grouped as ST_other.

| K-locus | Invasive | Non-Invasive | STs<br>(n=count) | Total |
| --- | --- | --- | --- | --- |
| KL64 | 122 | 152 | ST231 (n=9), ST147 (n=102), ST395 (n=90), ST14 (n=6), ST_other (n=13), ST2096 (n=54) | 274 |
| KL51 | 121 | 128 | ST231 (n=212), ST147 (n=20), ST_other (n=4), ST16 (n=13) | 249 |
| KL2 | 46 | 42 | ST14 (n=55), ST_other (n=17), ST101 (n=2), ST11 (n=5), ST15 (n=9) | 88 |
| KL10 | 16 | 34 | ST147 (n=46), ST_other (n=4) | 50 |
| KL36 | 22 | 20 | ST_other (n=1), ST437 (n=41) | 42 |
| KL81 | 17 | 17 | ST147 (n=2), ST_other (n=5), ST11 (n=1), ST16 (n=26) | 34 |
| KL17 | 8 | 14 | ST_other (n=2), ST101-1LV (n=11), ST101 (n=9) | 22 |
| KL112 | 8 | 13 | ST15 (n=21) | 21 |
| KL52 | 8 | 9 | ST_other (n=1), ST38 (n=11), ST437 (n=5) | 17 |
| KL62 | 10 | 7 | ST_other (n=6), ST48 (n=11) | 17 |
| KL102 | 8 | 7 | ST_other (n=3), ST307 (n=12) | 15 |
| KL Other<br>(count <15) | 113 | 130 | ST231 (n=4), ST147 (n=9), ST395 (n=4), ST14 (n=1), ST_other (n=155), ST101 (n=1), ST1710 (n=12), ST23 (n=12), ST469 (n=12), ST307 (n=1), ST11 (n=13), ST15 (n=8), ST16 (n=11) | 243 |

**Table 2.**
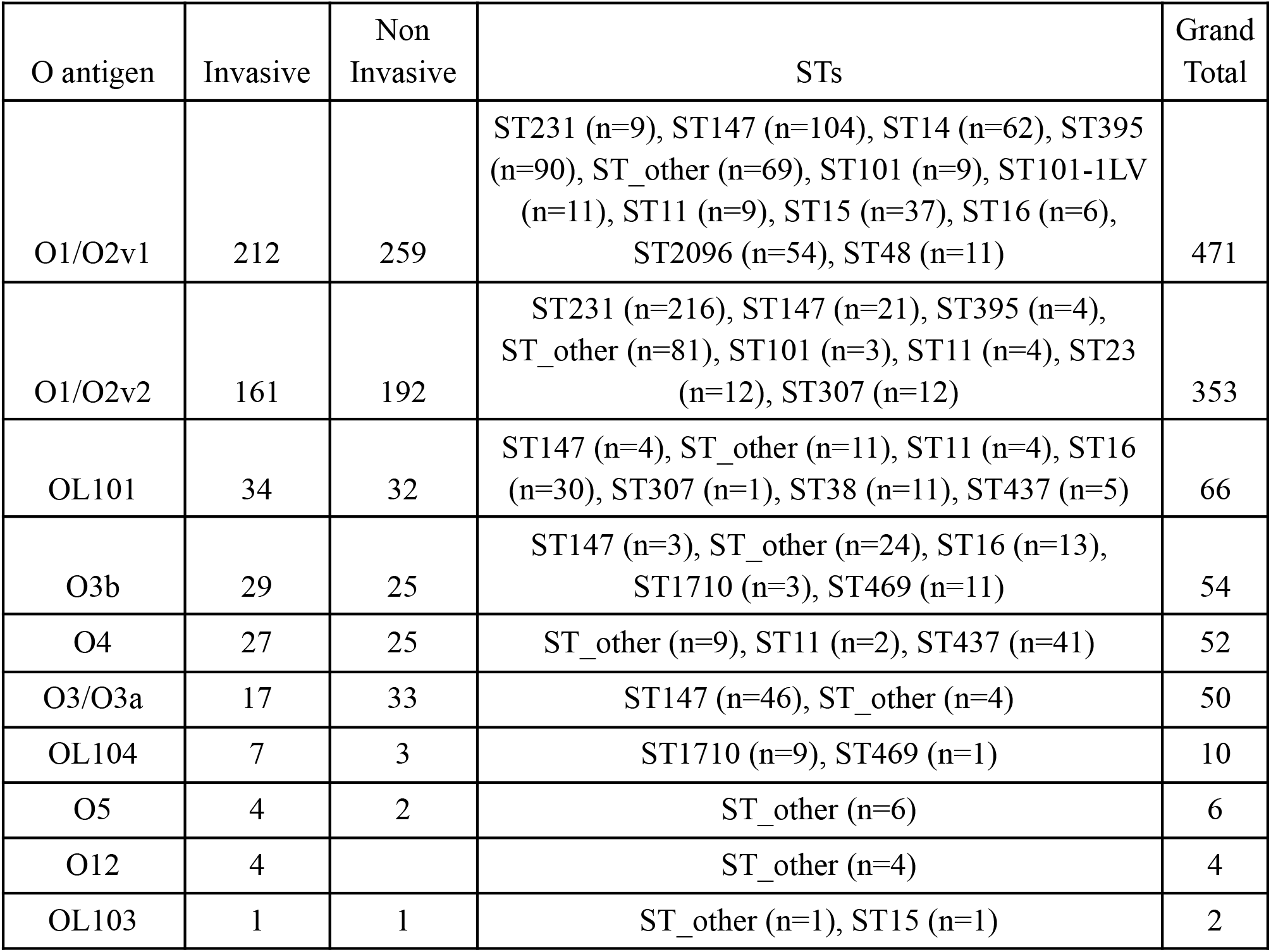
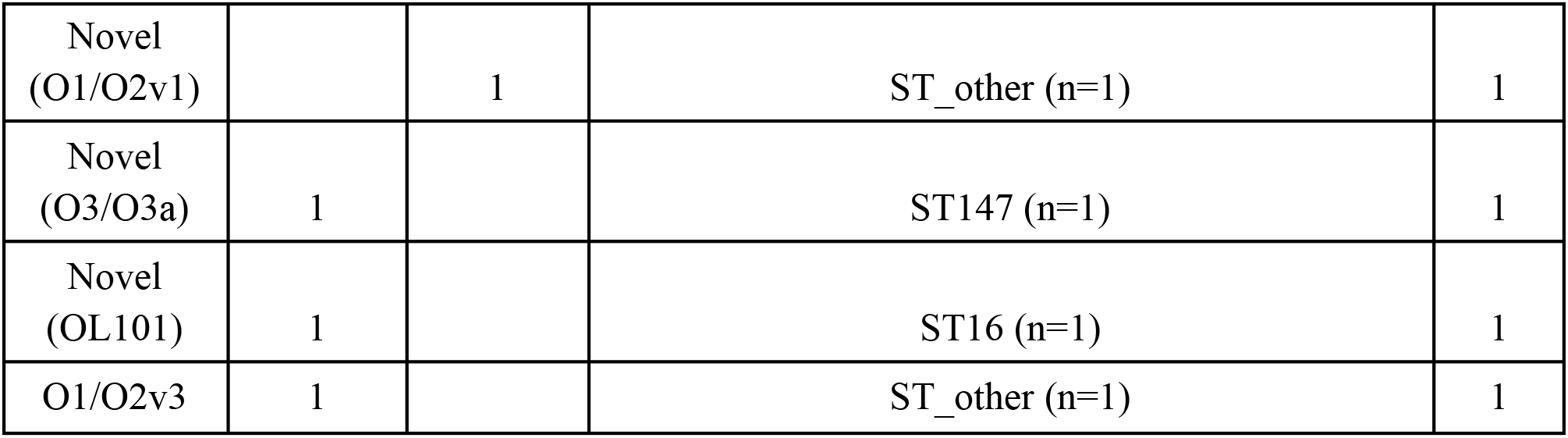
O-loci distribution among the invasive and non-invasive isolates and their associated STs. The top 10 STs are shown individually and others are grouped as ST_other.

**Figure 2.**
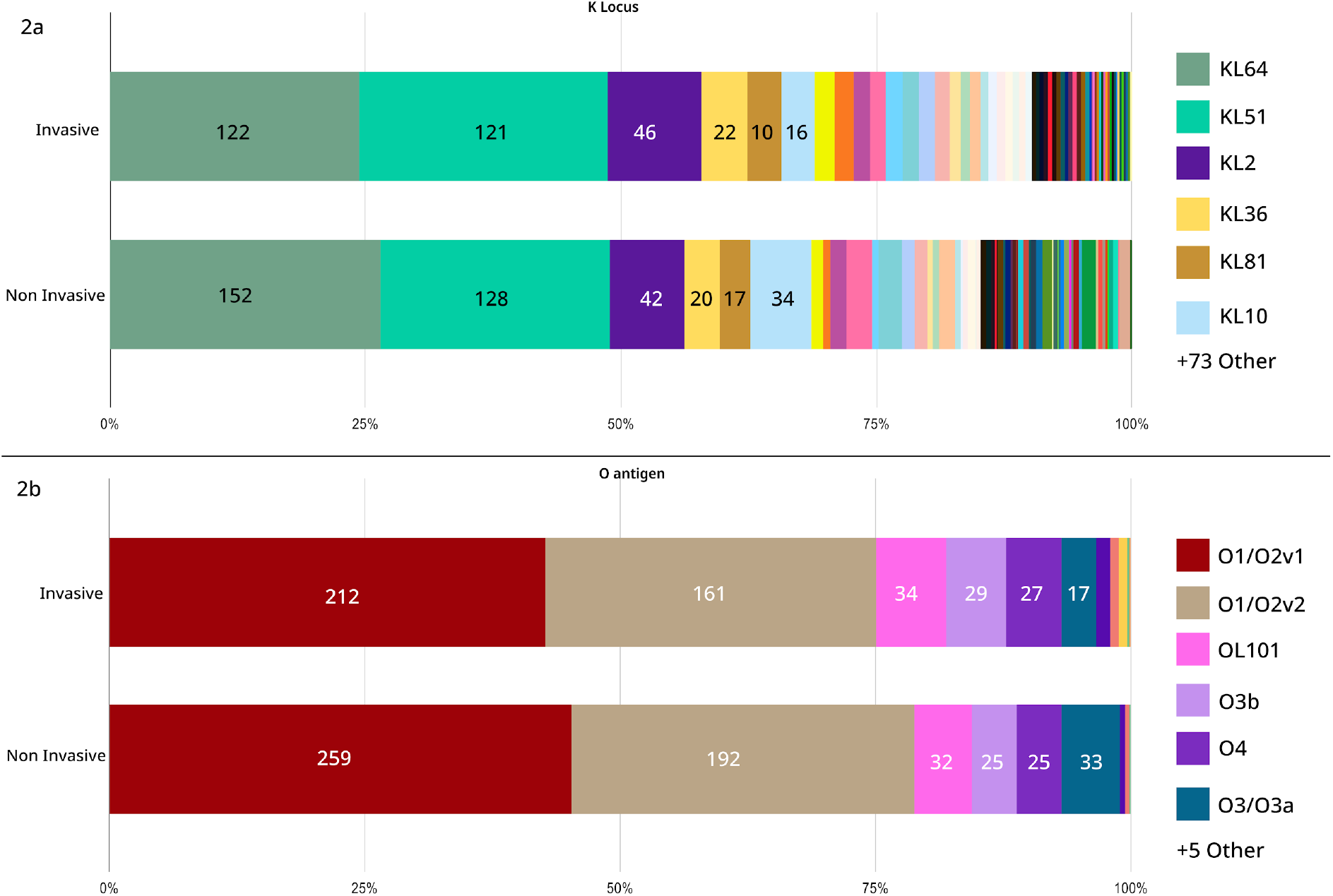
Distribution of KL and O types among invasive and noninvasive specimens. The bars are stratified by the number of samples in each KL and O type. The bars are stacked to 100%.

The theoretical coverage provided by multi-valent vaccines targeting increasing numbers of KL (ordered by KL frequency in the population) is shown in Fig. 3a. The diversity of KL types within our sample collection was assessed using Simpson’s Diversity Index (D). This index gives a numerical representation of the diversity of KL in the population, ranging from 0 (no diversity) to 1 (highest diversity). The KL were very diverse in age groups <2 years (D-value: 0.915), indicating that there is no specific KL associated with paediatric infections. The diversity among other age groups was also very high and the diversity calculation for each age group is provided in Suppl. Tables 3 and 4.

**Figure 3.**
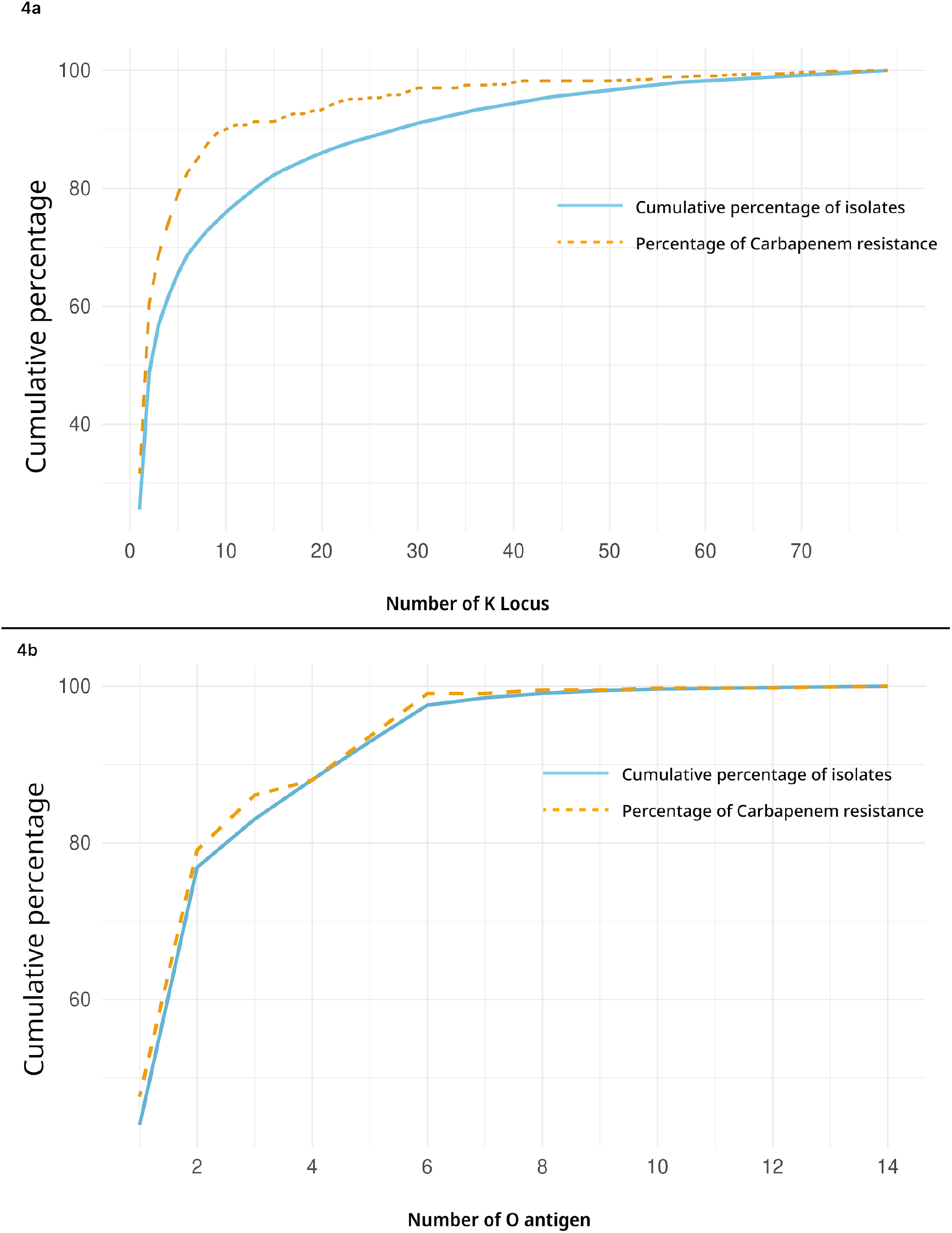
The cumulative coverage of the KL and the O types represents the percentage of isolates covered. The KL and O types are sorted with the highest to lowest frequency. The orange dotted line shows cumulative carbapenemase-positive strains.

Our study assessed the prevalence and associations of different O types with disease. Our collection had a diverse distribution of O antigens with 14 different O types. The most prevalent O-loci were O1/O2v1 (44%, 471/1072) and O1/O2v2 (33%, 353/1072), O10 (7%, 66/1072), which were collectively the top 3 O-loci constituting 83% (890/1072) of the collection. Other O-loci, such as OL101 (66/1072), O3b (54/1072), O4 (52/1072), and O3/O3a (50/1072), were also identified in Table 2). In the assessment of associations with sample sources, a Fisher’s exact test [42]) for O1/O2v1 (P-value: 0.388), O1/O2v2 (P-value: 0.696), and O101 (P-value: 0.445) revealed no significant associations, implying a similar distribution between invasive and non-invasive isolates. Serotype distributions in KL and O types showed no significant differences by age group, as confirmed by Fisher’s exact test. The cumulative coverage of O types shows that if 5 types are incorporated in a vaccine formulation, it will cover 90% of the isolates from different specimen types and disease conditions (Figure 3).

### Sequence types and their serotypes association

The 1072 genomes of *K. pneumoniae* encompass 119 different STs, some persisting for several years. Most of them have been reported globally while others are mainly restricted to Southeast Asia, such as ST231 (n=228), ST147 (n= 180), ST395 (n=94) and ST14 (n=62) (Supplementary Table 1). Analysis of the KL types revealed a distinct pattern of variation, where some STs were associated with single K-locus types and others had multiple K-locus types (Figure 4). For example, within ST231, KL51 is the dominant allele while KL64 is present in a small subset. The second most prevalent ST, ST147, also exhibited remarkable K-locus heterogeneity, hosting KL64 (n=110), KL51 (n=21), and KL10 (n=49) alleles and three additional K-loci, highlighting the intricate diversity within this clonal group. On the other hand, ST395 exclusively harboured KL64, while ST14 exhibited a strong association with KL2. This suggests lineage-specific selective pressures or functional constraints shaping KL diversification within the bacterial population.

**Figure 4.**
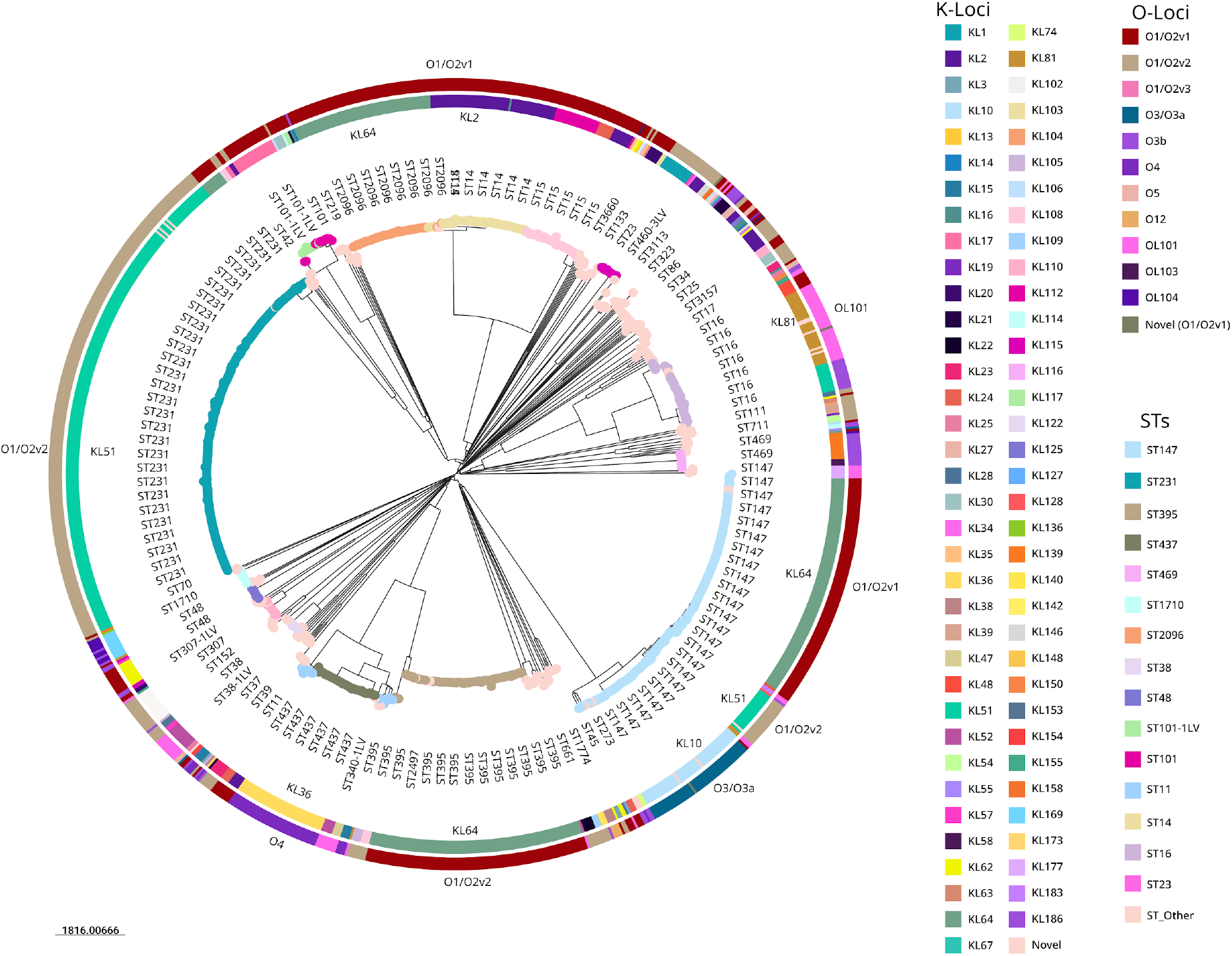
The circular view of the SNP tree was constructed using 100 bootstrap replicates. The tree is midpoint rooted and the scale bar represents the SNPs per variable site. The inner ring represents the K-loci types and the outer ring represents the O antigen types. The nodes are coloured by ST.

O serotypes were strongly linked with STs. O1/O2v2 was seen in 220 isolates of ST231 and 21 of the ST147 isolates but ST147 also carried clade-specific associations with O1/O2v1 and O3/O3a. ST395, ST14, ST15, ST2096 and ST101 only carried O1/O2V1. ST16 was associated with 3 different O-types namely O101, O1/O2v1 and O3b. Other minor STs had varied O-types but some were associated with single O-types. This pattern of O-types suggests ancient independent acquisition of O-types with subsequent clonal spread. The distribution of K- and O-loci among different STs is shown in Figure 4 and also in the microreact views (microreact.org/klocusvsst and microreact.org/olocusvsst)

### Geographical distribution of KL and O antigens across India

We also examined the geographical distribution of KL and O antigens across India, revealing the diversity of serotype prevalence. The distribution of KL and O antigens was heterogeneous, with some loci being more prevalent in particular parts of the nation than others. The ST147 clone with KL64 and O1/O2V1 locus and the ST395 clone with KL64 and O1/O2V1 locus were the two most prevalent types in the Northern part of the subcontinent. Twelve isolates of ST1710 having KL169 and O104 or O3B were seen in only one sentinel site from Gujarat.

From the Simpson’s diversity index, the calculated overall D value of 0.8663956 indicates a relatively high level of diversity among KL types. On the other hand, the regional variation in the K loci diversity is quite remarkable. The Suppl. Table 5 reveals the Simpson’s Diversity Index for six major political zones across India. We observed that specific zones exhibited a higher degree of diversity in KL types compared to the others. Notably, the Southern and Central zones displayed the highest diversity in K loci, with Simpson’s Diversity Index values of 0.914 and 0.894, respectively. The diversity was relatively lower in the Northern and Western zones (0.759 and 0.794, respectively) (Suppl. Table 5). The major KL types KL64 and KL51 and O types O1/O2v1 and O1/2v2 were distributed across all the zones and the sentinel sites.

### Virulence factors associated with capsular (KL) and O antigen serotypes

We screened for all 6 known virulence factors in *K. pneumoniae*, i.e. yersiniabactin (*ybt*), aerobactin (*iuc*), salmochelin (*iro*), the genotoxin colibactin (*clb*), and the hyper mucoid locus *rmp*ADC and *rmp*A2. From the Kleborate virulence scoring, 314 out of 1072 (26%) isolates in our collection had a virulence score of 4. The most common KL found in these was KL51 (179/314, 57%) including 132 ST231, followed by KL64 (91/314, 29%) including 38 ST2096. Only 9/14 isolates of KL1, a globally known hypervirulent serotype, had a virulence score of 5 with all the virulence factors. 69/88 isolates of KL2 had a virulence score of 1 and only 9/88 of them carried the hypervirulence marker aerobactin with a virulence score of 4. There was no specific association of virulence factors with the KL or O antigen types (P>0.05). These results show no particular association between serotypes and virulence factors. These findings underscore the complex and varied nature of virulence among *K. pneumoniae* isolates, highlighting the need for further research to elucidate the factors influencing pathogenicity in this bacterium. Interestingly, it was observed that conventionally described hypervirulent clones did not carry virulence markers, while other regional serotypes exhibited the presence of hypervirulence markers, adding a layer of complexity to the understanding of virulence in *K. pneumoniae*.

### Resistance profiles associated with capsular (KL) and O antigen serotypes

In this study, Kleborate identified 64 distinct genes associated with antimicrobial resistance. These genes are linked to resistance across 10 antimicrobial classes. All isolates carried at least one genetic resistance determinant and 73% (786/1072) were MDR having resistance to more than 3 classes of drugs. Notably, a significant majority, comprising 78% (841/1072), exhibited specifically carbapenem-resistance markers. The majority of the isolates in our collection (70%, 748/1072) had a resistance score of 2, followed by 14% (158/1072) of the isolates having a resistance score of 1, notably 4% (43/1072) had the highest resistance score of 3 and only 11% (123/1072) had a resistance score of 0.

Forty (12%, n=129/1072) of the KL types did not carry any carbapenemase genes. The other 39 types carried at least one carbapenamase gene. The predominant KL types, KL51 and KL64, exhibited varied associations with different carbapenemase genes, with KL64 primarily linked to *bla*_OXA-48-like_ variants (particularly *bla*_OXA-181_ (63/274) and *bla*_OXA-232_ (124/274)), while KL51 strains carried predominantly *bla*_OXA-232_ (183/249), followed by co-carriage of *bla*_NDM_ and *bla*_OXA-232_ (38/249). The carbapenemase gene distribution among different KL types is provided in Suppl. Tables 6 and 7. There was no significant association between O antigen types and carbapenemase genes (P>0.05). The predominant O types, O1/O2v1 and O1/O2v2 carried both *bla*_OXA-48-like_ and *bla*_NDM_ genes. Four different O types comprising 1.4% (16/1072) of the isolates did not carry any carbapenemase genes (Supplementary table 4). The acquisition of the carbapenemase genes was independent of serotypes and largely driven by the STs. The cumulative distribution of the KL and O types in Figure 3 showed that the vaccine formulas against *K. pneumoniae* in India need to incorporate either ∼20 KL types to cover >85% of the carbapenemase-producing population or 4 OL types to cover the same population.

## Discussion

*K. pneumoniae* is a human commensal and opportunistic pathogen that can cause severe hospital-acquired infections, especially among patients with compromised immune systems. *K. pneumoniae* infections are tough to cure due to the organism’s thick capsule [43] and the emergence of MDR strains has made the majority of current antimicrobials ineffective [44]. Effective measures for preventing *K. pneumoniae* infections are desperately needed and vaccines are one of the proposed alternatives [18]. Capsular polysaccharides in *K. pneumoniae* have emerged as promising targets for vaccine development due to their role in virulence and the bacteria’s resistance mechanisms [19]. This study aimed to comprehensively investigate serotype prevalence and epidemiology in India, a region grappling with antimicrobial resistance and healthcare-associated infections.

The analysis of 1072 *K. pneumoniae* isolates from 2014 to 2022 in India revealed significant genomic diversity within the KL and O types. This highlights a varied genetic landscape among *K. pneumoniae* strains circulating in Indian hospitals. The extensive diversity observed within the KL of *K. pneumoniae* poses a considerable challenge for vaccine development. With the identification of 79 distinct KL types and the top three KL (KL51, KL64, KL2) collectively representing only 59% of the isolates, it is evident that creating a vaccine or monoclonal antibodies targeting a wide spectrum of prevalent KL types will be complex. The recently described development of anti-KL64 antibodies would only cover about 26% of the Indian *K. pneumoniae* isolates [45]. Several studies have consistently reported a high diversity of KL types in *K. pneumoniae* strains across different regions [46]; [47]; [48]; [28]; [49]. In Southeast (SE) Asia particularly, studies have highlighted the regional disparity of KL types in *K. pneumoniae* strains causing infection. For instance, a study by Holt and colleagues [7] found that KL2 and KL47 were predominant in Thailand, while KL107 was more prevalent in Cambodia. Similarly, a study by Wyres and colleagues [28] reported variations in KL types among *K. pneumoniae* isolates in Malaysia, with KL1, KL2, and KL102 being the most common. The development of the Inventprise 25-valent pneumococcal conjugate vaccine candidate, utilising the patented Hz-PEG-Hz linker technology platform, represents a significant advancement in vaccine technology [50]. It is anticipated that this vaccine candidate will offer the broadest coverage against pathogenic pneumococcal serotypes encountered by populations worldwide, irrespective of geographical location. Similarly, if a *K. pneumoniae* vaccine targeting 25 KL types were to be developed, it could potentially provide coverage for up to 90% of the Indian *K. pneumoniae* population along with 85% of CRKP clones, illustrating the profound impact such a vaccine could have on public health and disease prevention efforts.

Research has highlighted the variability in disease associations linked to specific KL types. Some studies have identified certain KL types, such as KL1 and KL2, as being more prevalent and associated with poorer disease outcomes [51]; [52]; [53], particularly in hypervirulent strains causing community-acquired invasive infections. Conversely, others have noted the rarity of these KL types associated with severe infections in certain geographic regions [54]; [27]; [49]. Our study also highlights that certain KL types within *K. pneumoniae* strains were found in both invasive and non-invasive diseases, indicating their versatile pathogenic potential. Additionally, we did not observe an association between hypervirulence and certain KL types or disease severity, as noted before. This finding further complicates our understanding of the complex relationships between *K. pneumoniae* virulence factors and disease manifestation, emphasising the need for further research to elucidate the mechanisms underlying KL-associated pathogenicity.

We discovered a considerable diversity of 14 different O types among *K. pneumoniae* isolates compared to previous studies [28]; [55]. However, a few strong O-loci, especially O1/O2v1, O1/O2v2, and O10 stood out, accounting for a significant proportion (83%) of the isolates analysed. The cumulative coverage analysis revealed that incorporating only five types of O antigens (O1, O2v1, O2v2, O10 and O3b) would cover approximately 90% of the isolates across different specimen types and disease conditions in India. Previous studies have also explored the inclusion of a narrower selection of O types in vaccine formulations. For instance, Wyres and colleagues [28] suggested that incorporating a subset of prevalent O antigens could provide significant coverage against *K. pneumoniae* infections. Studies have highlighted the importance of targeting specific O types, such as O1, O2, and O3b, in vaccine development strategies due to their high prevalence and association with disease manifestation [13]. A recent study reported a heptavalent O-antigen bioconjugate vaccine [56] which exhibits promising efficacy against some, but not all, *K. pneumoniae* isolates. While some studies [57] say that O antigen is accessible by antibodies irrespective of the capsule type other studies are highlighting that hyperproduction of CPS may inhibit the vaccine-induced O-antigen antibody binding [57]; [58]. A more recent study evaluating monoclonal antibodies against *K. pneumoniae* ST147_NDM-1_, concluded that highly bactericidal anti O-antigen antibodies are not protective against this hypervirulent and pan drug-resistant strain ([45]). This suggests more studies are needed to clarify the effectiveness of anti O-antigen vaccine or monoclonal antibody formulations against the broader *K. pneumoniae* population.

## Conclusion

The findings provide crucial insights into the genetic diversity, evolution, and potential adaptation mechanisms of K and O loci within the *K. pneumoniae* Indian population, with implications for understanding its epidemiology. Additionally, the insights regarding its diversity underscore the challenges in formulating vaccines or monoclonal antibodies that adequately cover the diverse array of *K. pneumoniae* strains including the carbapenemase-producing ones. The lack of a straightforward correlation between specific serotypes and virulence factors challenges conventional assumptions about hypervirulence clones and necessitates further exploration into the multifaceted nature of virulence determinants in *K. pneumoniae*. Understanding the epidemiology of *K. pneumoniae* can help tailor effective prophylactic and therapeutic solutions against KL and O antigens in India.

## Author contributions

This study was conceptualised by V.S. and D.A., V.S. performed the genomic analysis of the samples collected in this study. S.S. was involved in statistical analysis, table generation, and figure generation. N.C. guided the manuscript preparation and reviewed the manuscript. Funding for the study was provided through grants to K.L.R and D.A., G.N. and H.G.K. were involved in the sample collection. H.G.K. performed the microbiology part of the analysis & G.N. did the sequencing. All authors reviewed the manuscript and suggested improvements.

## Acknowledgements

Members of the NIHR Global Health Research Unit on Genomic Surveillance of Antimicrobial Resistance: Sophia David, Monica Abrudan, Julio Diaz Caballero, Emmanuelle Kumaran, Georgina Lewis-Woodhouse, Khalil Abudahab and Ben Pascoe of the Centre for Genomic Pathogen Surveillance, Big Data Institute, University of Oxford, Old Road Campus, Oxford, UK, and Wellcome Genome Campus, Hinxton, UK; Pilar Donado-Godoy of the Colombian Integrated Program for Antimicrobial Resistance Surveillance, Coipars, CI Tibaitatá, Corporación Colombiana de Investigación Agropecuaria (AGROSAVIA), Tibaitatá, Mosquera, Cundinamarca, Colombia; Shreedhanya D M, Shincy M. R., Sravani D., K. N. Ravishankar of the Central Research Laboratory, Kempegowda Institute of Medical Sciences, Bengaluru, India; Iruka N. Okeke, Anderson O. Oaikhena of the Department of Pharmaceutical Microbiology, Faculty of Pharmacy, University of Ibadan, Oyo State, Nigeria; Sonia Sia, Celia Carlos, Marietta L. Lagrada and June M. Gayeta of the Antimicrobial Resistance Surveillance Reference Laboratory, Research Institute for Tropical Medicine, Muntinlupa, the Philippines; John Stelling, The Brigham and Women’s Hospital, Boston, MA, USA; and Carolin Vegvari, Imperial College London, London, UK. NMAMIT, Dept. of Biotechnology: Ujwal P, Vidya S M and Chethan D M. GHRU India Consortium: Dr. Anuradha Sharma, AIIMS, Jodhpur, Rajasthan, India; Dr. Ujjwayini Ray, Apollo Multi Scpeciality Hospital, Kankurgachi, Kolkata, West Bengal, India; Dr. Manick Das, Apollo Medical College, Hyderabad, India; Dr. Maneesha Sahu, BALCO Medical Center, Raipur, Chhattisgarh, India; Dr. Aruna Poojary, Breach Candy Hospitals, Mumbai, Maharashtra, India; Dr. Lakshmi, Biocare Research Lab, Gandhinagar, Gujarat, India; Dr. Shwetha, Bangalore Medical College, Bangalore, Karnataka, India; Dr.malini sahroff, BPS womens Medical college, Sonipat, Haryana, India; Dr. Shubranshu mandal, Calcutta Medical Research Institute, Kolkata, West Bengal, India; Dr. Frincy, Excel Health care, Guwahati, Assam, India; Dr. Anitha, Government medical college, Trichy, Tamil Nadu, India; Dr. Varsha Guptha, GMC, Chandigarh, India; Dr. Namrata Rai, Indira Gandhi Institute of Medical Sciences, Patna, Bihar, India; Dr. Bhattacharyya, All India Institute of Hygiene & Public Health, Kolkata, West Bengal, India; Dr. Naveena, Sri Jayadeva Institute of Cardiovascular Sciences and Research, Bangalore, Karnataka, India; Dr. Sujatha, Jawaharlal Institute of Postgraduate Medical Education and Research, Pondicherry, Pondicherry, India; Dr. Sheetal Verma, King George’s Medical University, Lucknow, Uttar Pradesh, India; Dr. Shrikala baliga, Kasturba Medical College, Mangalore, Karnataka, India; Dr. Lakshmi, Kamineni Hospital, Hyderabad, Telangana, India; Dr. Shalab Malik, Dr Lal PathLabs, Delhi, India; Dr. Vishwanath, Mediquest Diagnostics, Hyderabad, Telangana, India; Dr. Sohan Lal, Malabar Institute of Medical Sciences, Kozhikode, Kerala, India; Dr. veenakumari, NIMHANS, Bangalore, Karnataka, India; Dr. Milan, Neugen Laboratories, Rajkot, Gujarat, India; Dr. Venkatraman Kandi, Prathima Institute of Medical sciences, Karimnagar, Telangana, India; Dr. Smita Sood, Rukmani Birla Hospital, Jaipur, Rajasthan, India; Dr. Keerthi Lakshmi, RajaRajeswari Medical College, Bangalore, Karnataka, India; Dr. Jyothi EK, SCTIMST, Thiruvananthapuram, Kerala, India; Purna chandra, Kolar medical college, Kolar, Karnataka, India; Dr. Vaidehi, Sundaram Medical Foundation, Chennai, Tamil Nadu, India; Dr. Jagatheeswary, Saveetha Medical College, Kuthambakkam, Tamil Nadu, India; Dr. B G Vishwanath, Sreekar Lab, Hyderabad, Telangana, India; Dr.Shruthi Uppoor, Siddiah Referral Hospital, Bangalore, Karnataka, India; Dr. chitra rajalakshmi, Trichy SRM Medical College Hospital and Research Centre, Trichy, Tamil Nadu, India; Dr. Mohit Bhatia, AIIMS, Rishikesh, Uttarakhand, India; Dr. Sowmusharee, VIMSAR, Burla, Odisha, India; Dr. Malini Sahriff, Vallabhbhai Patel Chest Institute, Delhi, India

