## Supplementary material 2 for "Geographical distribution, disease association and diversity of *Klebsiella pneumoniae* KL and O antigens in India: roadmap for vaccine development"

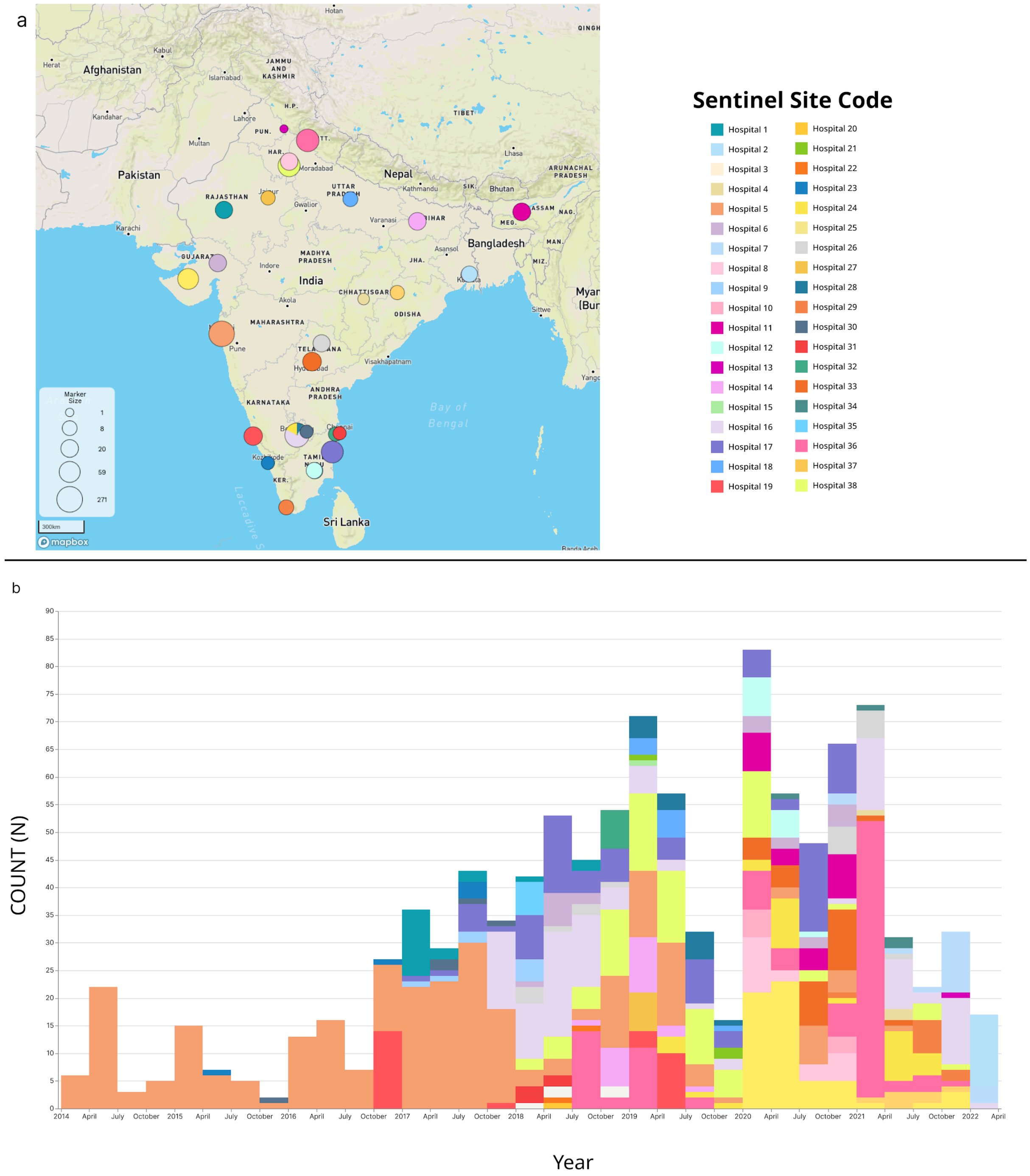


**Supplementary Figure 1A:** Geographical distribution of the 1072 *K. pneumoniae* isolates collected across 38 different sentinel sites in India.

**Supplementary Figure 1B:** Timeline of the samples collected from the year 2013 to 2022 for this study.
